# Hippocampal tangential insertions of high-density silicon probes in head-fixed mice enhance spatial sampling

**DOI:** 10.64898/2026.01.22.701007

**Authors:** Annu Kala, Vera Evander, John J. Tukker, Jens Kremkow, Dietmar Schmitz, Jérémie Sibille

## Abstract

The hippocampus is a key structure within the medial temporal lobe which plays a central role in the formation and consolidation of declarative memories. Despite technological progress, a full understanding of hippocampal rhythms, neuronal population activity, and the underlying connectivity remains elusive, partly due to its complex morphology in a croissant-like shape which limits coronal and sagittal approaches. To address this limitation, we propose two different tangential insertions of high-density electrode probes (Neuropixels 1.0) aligned to either the dorsal or ventral sections of the hippocampus. Each approach enables the placement of up to 384 recording sites within the hippocampal formation. Both insertions are performed in head-fixed conditions, which can be easily combined with detailed monitoring of the animals’ behavioral states including spontaneous running and facial movements. The two proposed tangential dorsal and ventral insertions capture almost continuously the physiology of the hippocampus across 3.84 mm of tissue. We show that when placed across the pyramidal layer, such insertions can harness ripple oscillations from approximately 130 channels, corresponding to about ∼1.3 mm of tissue, with a coverage of 2 channels every 20 µm. Furthermore, such insertions enabled us to record up to 400 hippocampal neurons simultaneously from a single probe, endowing us with the capability to capture putative functionally connected neuronal pairs. Although the conventional orthogonal/vertical approach is optimal for recording the *laminar* organization of hippocampal activity, we demonstrate how, for the *longitudinal* axis of hippocampus, tangential insertions can substantially increase spatial reach and sampling resolution, enabling a more detailed description of activity along this axis.

## Introduction

The hippocampus is one of the most studied regions in the mammalian brain. In fact, the very first “unit” recordings from the mammalian brain were made with glass microelectrodes from hippocampal pyramidal cells in the cat (Renshaw et al., 1940). This was expanded to single-cell recordings from awake behaving rats, revealing the existence of place cells encoding the spatial location of the animal (O’Keefe & Dostrovsky, 1971). The introduction of techniques combining several wire electrodes (McNaughton et al., 1983; O’Keefe & Recce, 1993; Wilson & McNaughton, 1993) or multiple recording points arranged in an array (Buzsáki, 2004; Wise & Najafi, 1991) greatly expanded our knowledge of neural coding by enabling the recording of spatially distributed ensembles of cells. The most recent step in this development has been the introduction of high-density recording probes such as the Neuropixels probe (NP) (Jun et al., 2017; Steinmetz et al., 2021; Ye et al., 2025) with a denser spatial coverage enabling the recording of hundreds of single-units simultaneously. Higher numbers of recorded single-units not only enable the identification of relatively rare subpopulations but also maximize the identification of mono-synaptically connected pairs of neurons (Barthó et al., 2004; J. Csicsvari et al., 1998; English et al., 2017; Fink et al., 2025; Perkel et al., 1967; Tanaka, 1983) and high-dimensional descriptions of neural population code (Gardner et al., 2021; Nagelhus et al., 2023; Zutshi et al., 2025). For such recordings, the spatial layout of recording sites is of crucial importance, not only to guarantee the largest possible number of recorded neurons, but also to reveal possible spatial structure in their activity. In the hippocampus, as in other cortical tissues, such structure may be found in a laminar or sublaminar organization, which has been studied extensively using vertical insertions, but also across sub-regions (Centofante et al., 2023; S. Deadwyler et al., 1996; S. A. Deadwyler & Hampson, 1999; Fanselow & Dong, 2010; Hampson et al., 1996, 1999; Henriksen et al., 2010; Igarashi et al., 2014; Soltesz & Losonczy, 2018), where differences have been shown in connectivity and molecular expression profiles (Cembrowski et al., 2016; Kaulich et al., 2025; Lein et al., 2004; Thompson et al., 2008; M. Wang et al., 2016; Yao et al., 2021).

Similarly, local field potential (LFP) recordings in the hippocampus have been studied for a long time (Jung & Kornmüller, 1938), and oscillatory activity in various frequency ranges has been closely implicated in hippocampal function (Royer et al., 2010; van Bree et al., 2025; Vanderwolf, 1969). Studies using vertical approaches showed that these oscillations are organized in a strongly laminar manner, with specific “depth profiles” (Berényi et al., 2014; Brankack et al., 1993; Buzsáki et al., 1983; Castelli et al., 2025; Csicsvari et al., 1999; Green et al., 1961; Lasztóczi & Klausberger, 2014; 2016; Lopes-Dos-Santos et al., 2025; Paleologos et al., 2025; Winson, 1976). In addition, the use of multiple vertical probes in different locations of the hippocampus has revealed propagation patterns of theta waves (∼6-12 Hz; (Jackson et al., 2014; Lubenov & Siapas, 2009; Patel et al., 2012) and sharp-wave associated ripples (SWRs; ∼130-250 Hz) within and across subregions. Although these approaches have produced an incredible amount of data and insights, they do have a limited spatial resolution in terms of number of recording sites in “non-vertical” orientations, particularly along the antero-posterior axis of the hippocampus.

Finally, recording the brain activity from rodents in head-fixed conditions has supported a wealth of discoveries over the last two decades (Harvey et al., 2009; Niell & Stryker, 2010; Saleem et al., 2018; Siegle et al., 2021; Steinmetz et al., 2021; Ye et al., 2025). The correspondence between brain activity in freely moving and head-fixed running animals is still debated (Minderer et al., 2016); nevertheless, accumulated evidence suggests that in head-fixed recordings (combined with a virtual reality system) both the visual cortex (Saleem et al., 2018) and the hippocampus (Harvey et al., 2009) exhibit visual and spatial modulations. Furthermore, head fixed conditions permit precise control of different types of sensory cues (Bourboulou et al., 2019) and even enable single cell patch-clamp in the hippocampus (Bittner et al., 2015; Harvey et al., 2009), which revealed a new form of behaviorally driven plasticity (Bittner et al., 2017). Although head-fixation limits the study of certain aspects of behavior (Aghajan et al., 2015; Ravassard et al., 2013), it allows the characterization of more precise movement-related brain activity along with highly complex multi-probe insertion(s) at intricate angle(s).

Here, we present an alternative to vertical insertions of multiple wires or probe shanks. By inserting a single NP probe tangentially along the antero-posterior axis (Fig. 1, in either the ventral or dorsal parts of the hippocampus), we can substantially increase the number of channels (Fig. 1) within the hippocampus. This not only lets us map the distribution of LFP patterns at a high resolution across a large spatial extent (Fig. 2), but also allows us to capture units (Fig. 3) from an extended spatial domain across the pyramidal layer(s). Applying this approach in head-fixed recordings with high-resolution behavioral tracking makes it possible to correlate hippocampal activity with facial movements (Fig. 4). Furthermore, large numbers of functional connections can also be accessed along the longitudinal axis (Fig. 5). Overall, our approach enables the study of behavioral correlations with neuronal activity along the longitudinal axis of the hippocampus with higher precision.

**Figure 1:**
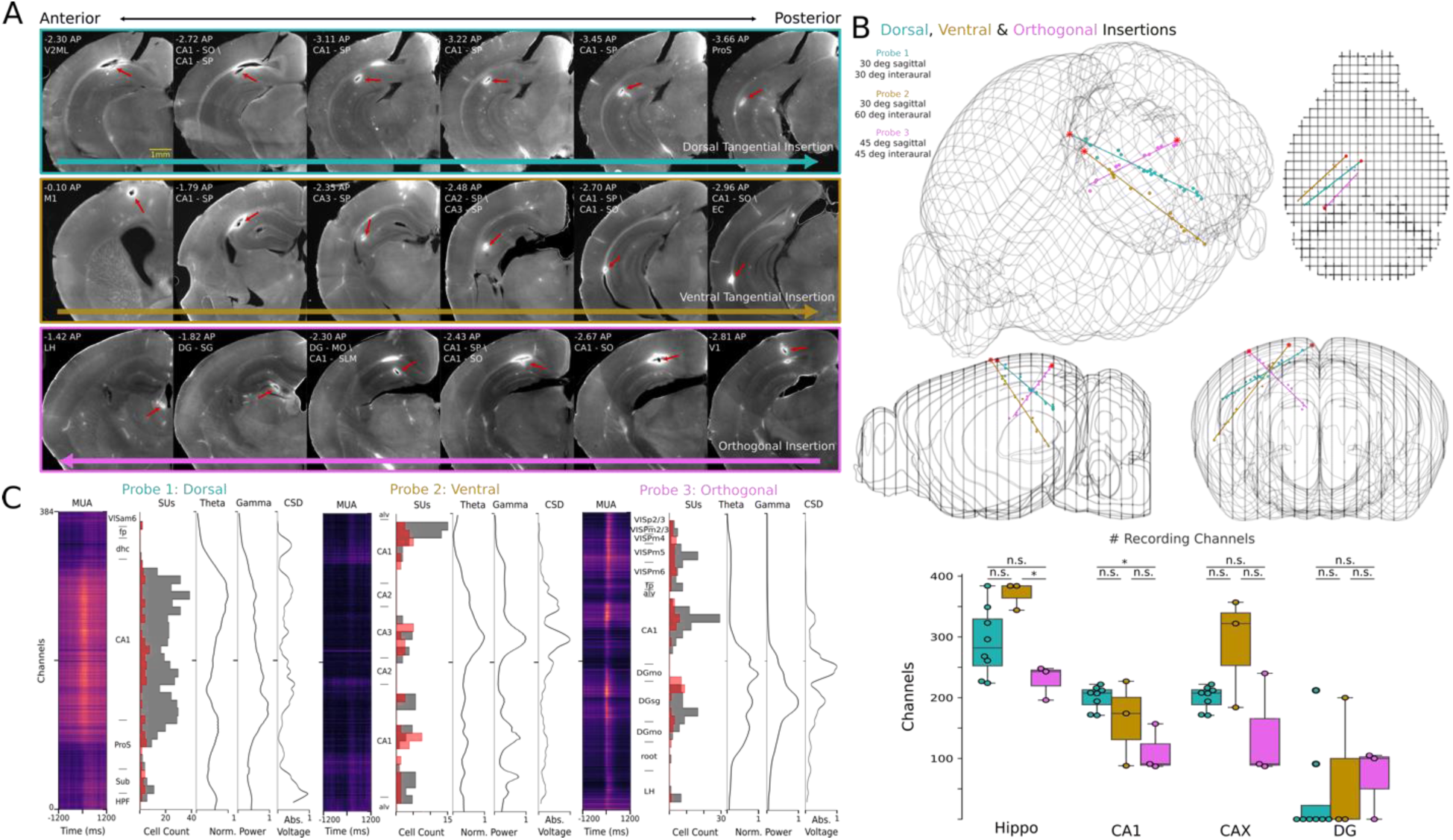
Hippocampal dorsal and ventral tangential insertions. **A.** Coronal sections from 3 different mice, illustrating the dorsal (top) and ventral (middle) tangential insertions, as well as a more standard orthogonal insertion (bottom), ordered in anterior (left) to posterior (right) planes. For each image, the AP distance relative to Bregma is indicated, along with the anatomical areas in which the probe track (red arrows) is located in this image. Colored arrows at the bottom of each row indicate the direction of each insertion. **B.** 3D reconstructions of the positioning of the probes in the left-brain hemisphere showing the probe placement for the three insertions (red dots indicate insertion points at the brain surface). 3D perspective (top-left), with corresponding sagittal (bottom-left), coronal (bottom-right), and interaural (top-right) projections. Below is the quantification of the number of recording channels in the hippocampus, CA1, general cornu ammonis regions (CAX), and DG by insertion(boxplot, bottom right). **C.** Illustration of the physiological alignment in a few recordings: for the dorsal (left), ventral (middle), and orthogonal (right) insertions also shown in B. For each of the 3 probes, for all recorded channels, plots show the mean normalized firing rate of multi-unit activity (MUA) aligned to movement onset (left), the number of isolated single-units (SUs/10 channels) obtained after spike sorting and curation (second from left; narrow-waveform units, red; broad-waveform units, grey), the LFP power in the theta (third from left; 4-10 Hz) and gamma (fourth from left; 30-100 Hz) bands, together with the channels’ anatomical locations deduced from the 3D reconstructions (between MUA and SU density). Plots also show the absolute value of the CSD along the probe when time-locked to theta trough (right) (cf. Sup. Fig. 2). The list of anatomical abbreviations can be found in supplementary table 2.

**Figure 2:**
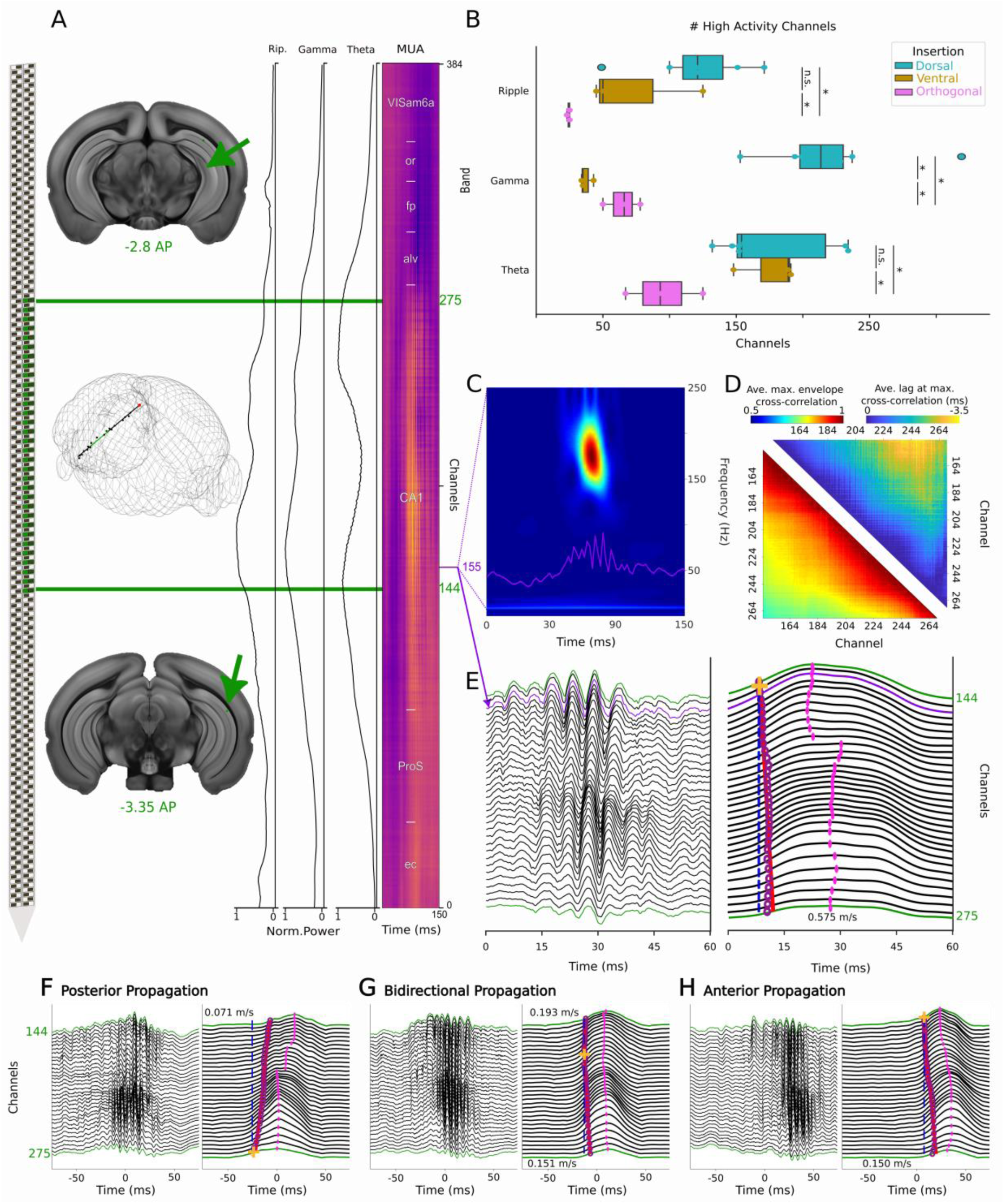
Bidirectional awake ripple propagation: **A.** Reconstructions of the example probe used throughout this figure. Green channels indicate the channels located close to the pyramidal layer shown in C-H (Left). Coronal slices and the green shaded section show the 1.29 mm region of dorsal CA1 along which individual ripples are tracked (Middle). LFP power profile in the Theta (4-10 Hz), Gamma (30-100 Hz) and Ripple (110-240 Hz) ranges, as in Fig. 1 (Right). Multi-unit activity in each channel overlayed with anatomical locations of the probe, aligned to the ripple event shown in C, E. See Sup. Table 2 for the list of abbreviations. **B.** Number of channels which have a normalized power greater than 0.5 for the theta and gamma frequency bands as indicated, with recordings aligned to movement onset as in figure 1C. Ripple quantification (top) indicates the number of channels along which ripple activity could be detected. **C.** Stockwell-transformed power spectrum of channel 155 centered on one ripple event. White trace is the corresponding LFP signal from this channel. D. Heatmaps showing the average max-envelope cross correlation (bottom) and the corresponding lag (top) between all channels within our ripple-area (channels 144-275, n = 105 ripple events). E. Raw LFP traces for an example ripple event (Left). Every four channels (green channels in A) are shown to remove putative issues from the NP1 checkboard pattern of neighboring channels (Right). Corresponding envelope of the Hilbert-transformed ripple-range-filtered signal for each channel. Yellow crosses indicate the channel in which the ripple was first detected. Purple circles indicate the time of crossing the 50% highest amplitude of the envelope. The red line is the linear polynomial fit across these time points measuring the different activation time, indirectly showing the propagation of ripples. Dashed blue lines represent zero lag w.r.t the yellow crosses. Average speed across the channels is calculated as (channels travelled*inter-channel-spacing)/(delay to 50% threshold crossing). Pink circles indicate the time of peak ripple power per channel. **F,G,H.** Same as D, but for events that begin and propagate in the different directions: posterior, anterior, and bi-directional.

**Figure 3:**
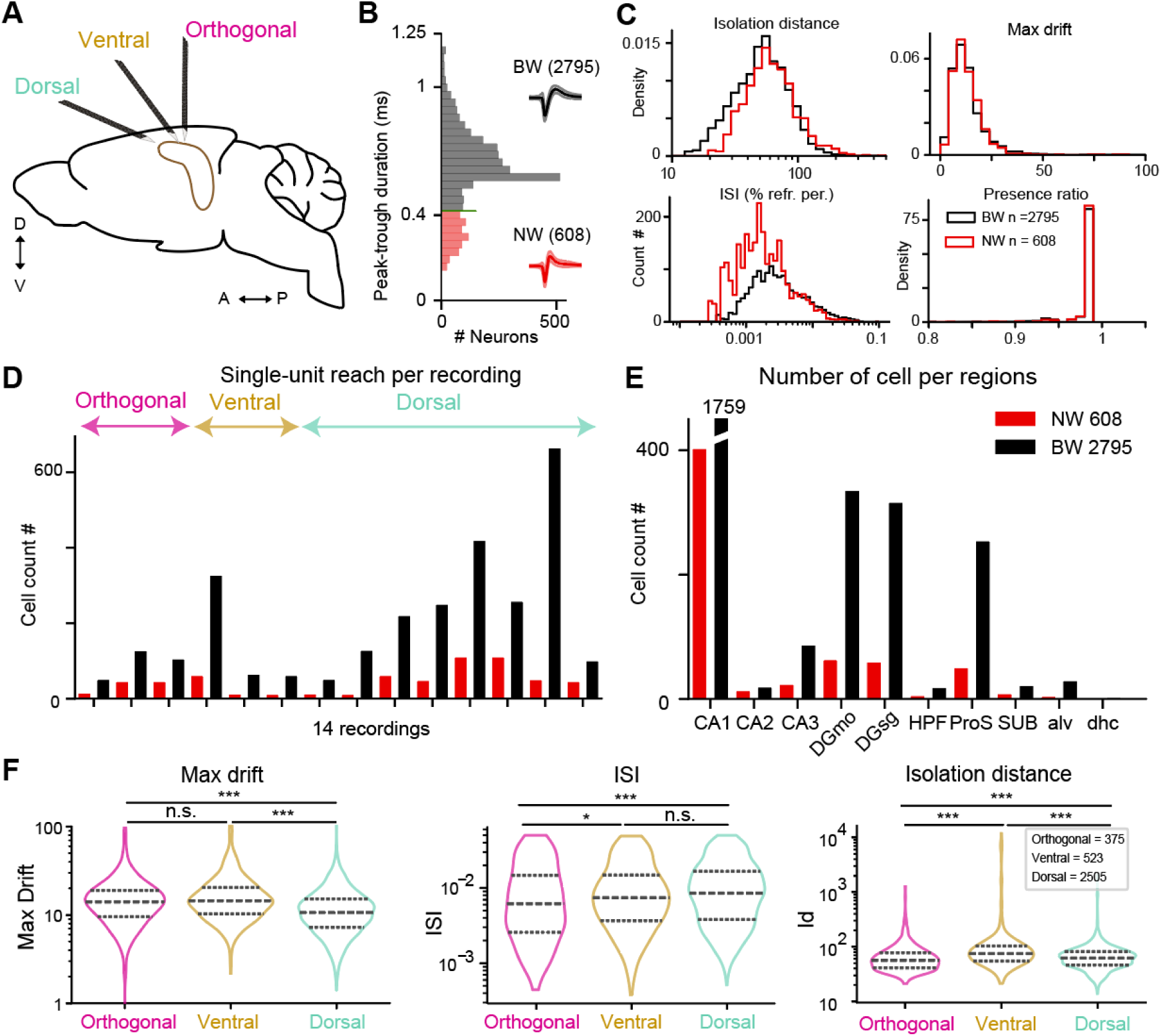
Single-unit reach, quality metrics: **A.** Illustration of the three insertion types. **B.** Quantification of the peak-trough duration illustrating the difference between broad and narrow waveforms (BW vs NW), in the hippocampal formation. **C.** Quality metrics of NW and BW neurons showing their high quality in isolation distance, inter-spike-interval (ISI), presence ratio, together with the higher stability (max drift) gained from the tangential insertion (n = 14 recordings from 14 mice). **D.** Quantification of NW and BW cell numbers for each insertion type. **E.** Quantification of NW and BW neuron numbers in each region identified from the 3D reconstruction (summed over all recordings). Note that the seemingly paradoxical higher number of neurons obtained in DGmo compared to DGstratum granulare may be related to the tangential positioning of the probe above the DG, thereby overestimating the number of channels in the DGmolecular layer when the spikes may stem from the DG-sg. **F.** Comparative quality metrics obtained in the different types of insertions, max drift (left), ISI, for refractory period violation (middle), and isolation distance (right). (n = 375/523/2505 neurons from n mice = 3/3/8 for orthogonal, ventral, and dorsal insertions respectively; Wilcoxon rank sum tests).

**Figure 4:**
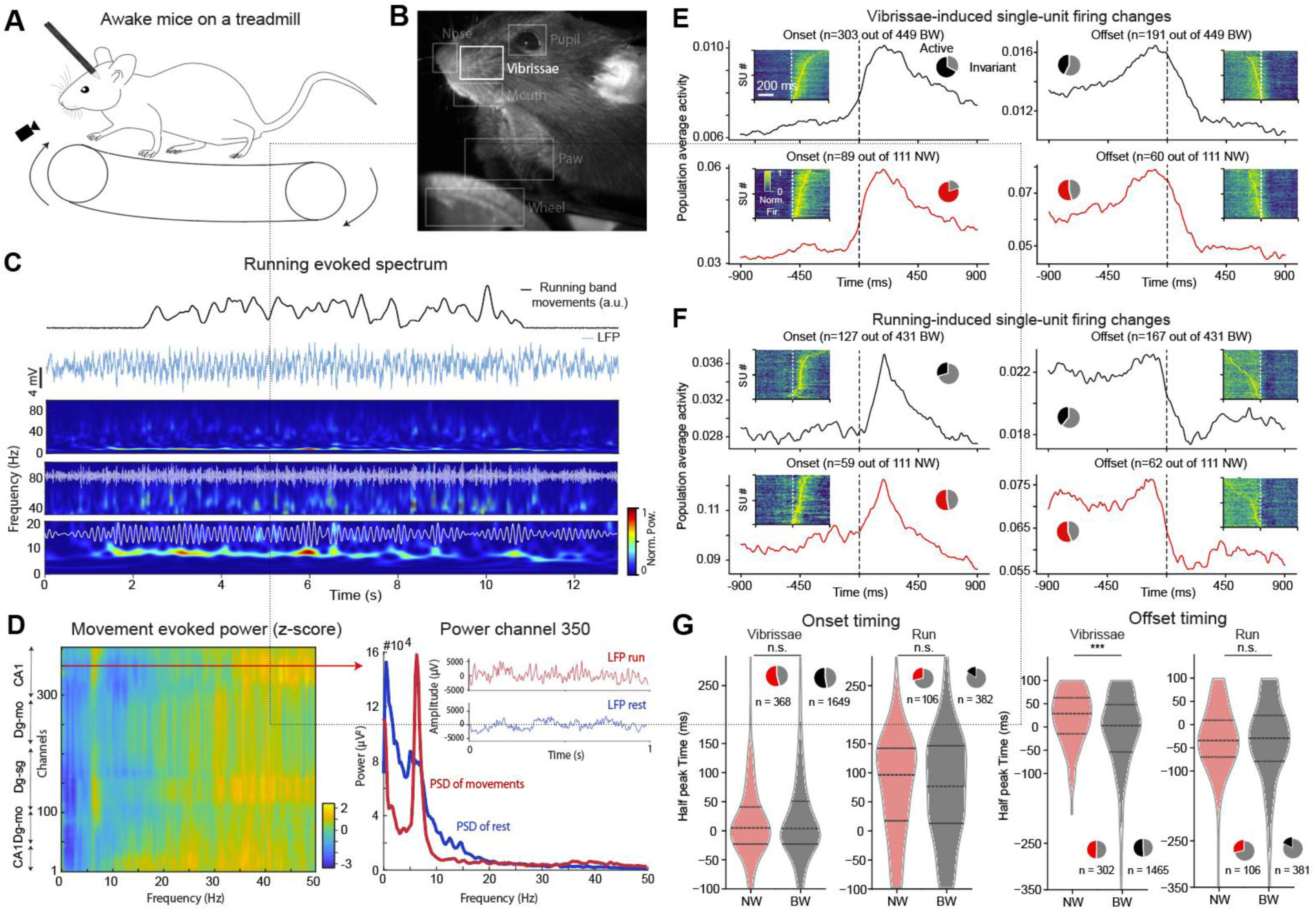
Coactivation of hippocampus with running and facial movement. **A.** Schematic of recording set-up: the acute head-fixed recording is combined with facial monitoring and quantification of the running behavior based on the analog signal measured from the running band. **B.** Illustration of the facial video, to show the region of interest used to characterize vibrissae movements (other regions labeled in light grey can be tracked; Sup. Fig. 6). Facemap was used to extract the singular value decomposition (SVD) to isolate moments of heightened activity versus rest (Sup. Fig. 6). **C.** Representative 13 seconds of the normalized running traces from the running band, in arbitrary units (Top). Corresponding LFP (Bottom). Time frequency representation from the LFP using Stockwell transformation (Top spectrogram). Zoomed high frequencies (40 – 90 Hz, Middle spectrogram). Zoomed low frequencies (0-20 Hz, Bottom spectrogram). The last two spectrograms also show corresponding bandpass-filtered LFP signal overlayed in white. **D.** Log-Normalized relative increase in power in each channel for movement periods vs silent periods (left). Blue and red lines show the corresponding PSD of movement and silent epochs in channel 350 (right). **E.** Mean population increase in firing of BW (top) and NW (bottom) neurons relative to the onset (left) and offset (right) of whisking moments (t = 0) within a single recording. Insets illustrate the normalized firing changes of all neurons showing a meaningful change of activity during onset and offset of facial movement. Pie plot insets illustrate the proportion of modulated (Active) versus invariant (Invariant) neurons during whisking (n = 1 mouse). **F.** Equivalent measures in the same recording of BW and NW neurons to the onset (left) and offset (right) of running bouts (n = 1 mouse). **G.** Dataset estimates of half-peak delay times of the hippocampal neuronal responses relative to the onset (left) and offset (right) of vibrissae and run movements. Violin plots show all the half peak times to illustrate the spread of the obtained values: the middle-dashed lines show the mean values; top and bottom lines show the 25^th^ and 75^th^ quartile. Pie plots show proportions of active (Active) versus invariant (Invariant) neurons throughout the dataset (n = 8 mice for running behavior, n = 14 mice for vibrissae movements; all statistics are done using the Wilcoxon rank sum test).

**Figure 5:**
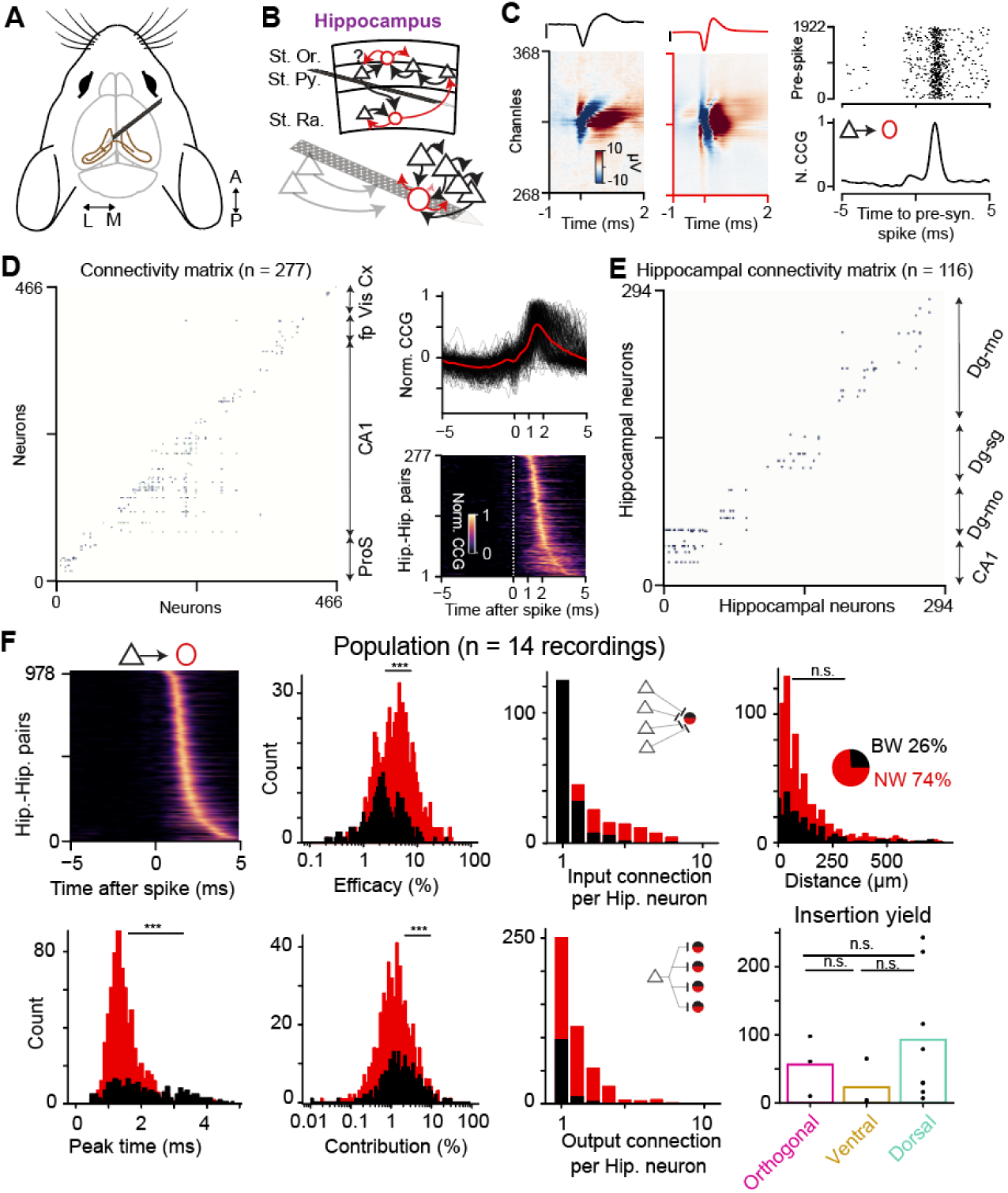
Functional excitatory connectivity within the hippocampus: **A.** Schematic of the NP placement for a tangential dorsal insertion. **B.** Schematic connectivity measure within the hippocampus between BW neurons to other neuron(s): other BW (black) or NW (red) neurons. **C.** Illustration of a single excitatory synaptic contact of a BW neuron toward a NW neuron. Each neuron is shown with its single channel waveform (top, scale bar is 50 µV) and below its 2D multi-channels waveform, illustrating its different spatial and temporal extent. Functional excitatory contacts are identified by their typical prominent peak at a monosynaptic time delay between the pre- and post-synaptic spiking activity (cf. raster plot and corresponding CCG, bottom right). **D.** Typical connectivity matrix in a single recording showing 277 excitatory connections between the available 466 neurons (neurons are organized based on their increasing channel position on the probe, left); with their corresponding reconstructed brain regions (double arrow middle). The corresponding normalized CCG profiles (top right; n = 277, 243 pairs are in the hippocampal formation) and the corresponding heat-map plotted signals (bottom right). **E.** Another connectivity matrix example, here illustrating more regional connections in this other recording. **F.** Summary and quantification of all available connections within the dataset (n = 980 pairs, from 14 recordings, n = 0-270 pairs per recording). A large majority of detected excitatory monosynaptic connections are toward NW neurons, as illustrated in the pie chart inset (22% of connections are BW to BW neurons). Quantifications of (top left to right): CCG profiles, efficacy, convergence, and distance between the connected neuronal pairs; (bottom left to right): peak times after spikes (with the mean population values), contribution, and divergence connectivity scheme. The last bottom right panel illustrates the differences of synaptic contact reach between orthogonal, tangential ventral and dorsal insertion (n = 3/3/

## 2. Materials and Methods

### 2.1 Animal surgery

All data presented were recorded in head-fixed conditions, with a protocol that was approved beforehand by the corresponding authority (Landesamt für Gesundheit und Soziales Berlin – G0298/18). Maximum care is dedicated to minimizing the number of animals used (n = 14 Adult C57BL/6J male mice from different litters) which all stem from local breeding facilities (Charité-Forschungseinrichtung für Experimentelle Medizin). A head post was implanted ∼ 3 weeks prior to the recording day. Such implantations were performed in a Narishige stereotactic frame using isoflurane (2.5 % in oxygen; Cp-Pharma G227L19A). Analgesia was delivered to the animal 1 day prior and 3 days post-surgical operations (Metamizol diluted in drinking water, 200 mg/kg body weight; Zentiva-Novaminsulfon). It is important to note that when the head-post was fixed, it was combined with a large dental cement-based crown (Paladur, Kuzler) to better fix the head-post on the skull, permitting a robust preparation that will stay over the 3-4 weeks of consequent manipulations. This crown was set up to grasp the lateral ridges of the skull and the frontal bone, to maximize stability. To facilitate dorsal tangential insertions which are relatively flat, it is important to reduce the crown height to 1-2 mm on the opposite side from the planned insertion. This crown was also made to surround the skull to hold fresh PBS during recording for grounding purposes. Mice were gradually habituated to head fixations on a running band (Phenosys©, Speedbelt 1601-0102, analog output 0-5 mV) for over three weeks, including breaks of a few days and monitoring of their wellbeing.

A second surgery was performed on the recording day to drill the craniotomy and locally remove the dura. A recovery period of at least 2 hours was given before recordings, and the wellness of the animals was assessed before initiating the recordings. The recordings were performed the same day as craniotomy to minimize the exposure of the tissue. Craniotomy, however, can also be performed a day prior to recording which improves the animal’s recovery from anesthesia. The craniotomy was performed with the following coordinates: -3.5 to -2.5 mm AP, 0 to +1 mm ML from Bregma for dorsal tangential insertions and -4 to - 3 mm AP, +2 to +3 mm ML from Bregma for ventral tangential insertions. The craniotomy for tangential dorsal insertion was purposely made larger to allow easier (re)insertion. During the craniotomy surgery, we used the drill to carve into the dental-cement crown and add a grounding wire, fixing it to the crown with extra dental cement, such that one end touched the skull within the crown while the other extended 3-5 cm in the air. The exposed grounding wire end was later used to connect to the NP external reference directly, which can be equipped with corresponding gold pins. For efficient noise reduction, the ground wire was immersed in the crown containing PBS. This ground wire replaced the classical screw in the skull above the cerebellum classically done for chronic implant. During the recordings, we prevented the bath from drying out by regularly refilling it with fresh PBS.

### 2.2 Insertion and staining

It is advised to prepare the micromanipulator’s placement beforehand (inserting with dummy probe for placement testing) to minimize the head-fixation time in the set-up during recording day. The micromanipulator arm (New Scale) was positioned on a custom Thorlabs mounting platform to achieve an overhead approach, which can also be done using a New Scale ring. The New Scale manipulator was positioned at an angle of 30° from the sagittal plane for both dorsal and ventral insertions. Using the rotating capabilities of New Scale arm, the moving head could be placed with the aligned probe either with a 30° angle from the interaural plane for dorsal insertions, or 60° for ventral insertions (Fig. 1 A & B, Sup. Fig. 1C). It is to be noted that our orthogonal insertions are performed at an angle of 45° from both the sagittal and the coronal plane (cf. Fig. 1), leading to higher unit yields compared to more standard purely vertical insertions (cf. Tab 1, Luo Brody 2020, Jun et al. 2017). Given the anatomy of hippocampus, angular insertion along the longitudinal axis encompasses relatively more cortical tissue compared to vertical ones, hence any deviation at the insertion point may amplify the errors in targeting the precise sub-structure of the hippocampus. Pre-recording practices, great attention to micromanipulator’s arm placement, attentive care on the skull flatness during head-post implant, and pilot brain slicing to confirm the probe placement will be considered as must-to-do prior to recording day. In addition, it is important to probe the possible variability between different mouse lines or ages to succeed with reproducible hippocampal tangential insertions. It is to be noticed that for tangential insertions, probes were inserted 5 to 6 mm deep in the tissue (particularly for ventral and dorsal insertion) to reach the full extent of the hippocampal formation, recording only from the lower shank of the NP. Once the probe is placed, the neuronal synchrony and LFP profile should be thoroughly checked to infer whether slight probes repositioning in different entry points of the craniotomy (Sup. Fig. 1C) would putatively improve the desired alignment.

On recording day, the tangential hippocampal insertions were initiated by slowly approaching the craniotomy and then moving carefully in the brain for at least 4-6 mm, followed by a withdrawal of 50-100 µm to release accumulated mechanical pressure. For the dorsal tangential insertions, given its flat angle, it is very important to avoid having any PBS in the crown in the beginning of the insertion in the tissue, otherwise the surface tension of the water will add extra bending of the probe, with an additional 10-15°, reducing reproducibility. It is therefore strongly advised to temporarily remove the grounding solution (PBS) at the time of contact of the probe with the brain tissue to guarantee an insertion aligned with the arm’s angles. Lastly, measuring Bregma and Lambda positions with the NP probe tip once the New Scale arm is in position can give extra measures for later positioning confirmation and ultimately better reproducibility.

As the probe is moved within the tissue, the presence of ripples and/or a large amount of spiking activity should be used to monitor the presence of CA1 or CA3 pyramidal layers along the channels (Fig. 1C & 2). This information may play an important role in assessing whether reinsertion at a different location, angle, or depth of the probe will be needed (Sup. Fig. 1). Importantly, given the proposed angle, the distance from the brain surface to CA1 can exceed 1.5 mm; therefore, a minimum of 5 mm of insertion should be done for ventral and dorsal tangential insertions (Fig. 1 and Sup. Fig. 1). Once a satisfying location was obtained, the probe was left to settle in the tissue for ∼30 minutes before starting the recording. The NP probe was coated with DiI (Abcam-ab 145311, in EtOH) before insertion. Once the recording was finished, the mice were sacrificed with a lethal injection of ketamine-xylazine (100 and 15 mg per kg body weight, respectively), before proceeding with cardiac perfusion via PBS and paraformaldehyde (PFA). Brains were left in PFA overnight, sliced coronally (100 µm) and mounted (DAPI-Fluoromount-G Biozol Cat. 0100-20). Mounted slices were pictured and the probe’s track was 3D reconstructed using SHARP-Track (Shamash et al., 2018) to relocate each single channel within the Allen Mouse Brain Common Coordinate Framework (Fig. 1)(Wang et al., 2020). Once the reconstruction was finished, only the channels, the cells, and the synaptic connections within the hippocampal formation (CA1-2-3 summarized as “CAX”, DGmo-sg-po summarized as “DG”, ProS, alv, dhc, HPF, SUB, cf. Fig. 1, Tab. 2) were kept for further analysis. Open-Ephys acquisition software (www.open-ephys.org) in combination with a PXIe system (National Instrument NI-PXIe-1071) was used to record NP probe signals (NP 1.0, IMEC).

### 2.3 Data acquisition and Local Field Potential extraction and coherence

LFP traces were recorded from the 384 channels of the Neuropixels probes (NP 1.0, sampled at 2.5 kHz). To analyze the prominence of different frequencies, power spectral density (PSD) was computed using Welch’s method with a window size of 4 s and 50% overlap. Power of theta, gamma, and ripple was obtained by summing the PSD within different frequency ranges (Theta: 4-10 Hz, Gamma: 30-100 Hz, Ripple: 110-240 Hz, Fig. 1C, 2A).

For a few analyzed dorsal tangential insertions (n = 6), theta and gamma power revealed large areas of the probes showing theta/gamma modulation during running movements (Fig. 4D, Sup. Fig. 4, cf. Rev. (Buzsáki, 2002)). To assess the spatial distribution of activity along the probes, we computed the current source density (CSD) for each recording by taking the second spatial derivative of averaged non-overlapping 1000 ms bouts of activity time-locked to theta-trough (Sup. Fig. 2). Because of the high spatial resolution we achieve within regions, a large channel separation (10 channels) was used in order to reduce the effects of volume conduction. To further probe the spatial distribution of theta activity, we computed PSD using FFT in a representative 20 s long running epoch across all the channels (Sup. Fig. 3). Summing the power in theta band (5-10 Hz) revealed a non-uniform pattern of theta activity across the probe (Sup. Fig. 3A, middle panel). Consistently, PSD traces from chosen channels (1-deep, 150-intermediate, and 300-superficial) (Sup. Fig. 3A, lower panel) show a similar gradient of theta activity. To visualize the full theta spectrum, we also constructed a frequency-by-channel power map to confirm the localization of theta prominence in the intermediate channels (Sup. Fig. 3B).

Using power spectrum extraction and its related normalized power, it became possible to compare the different profiles along the 384 channels from MUA, theta/gamma modulation, cell count, and anatomy (Fig. 1C). This alignment revealed that the probe can lie within the CA1 area and capture either the pyramidal layers, or the theta/gamma oscillations, or both, depending on its position below the pyramidal layers. As illustrated by other laboratories, theta/gamma oscillations are more prominent in the radiatum and lacunosum, compared to pyramidal layers where the ripple power is at its maximum (Paleologos et al. 2025). Coherent activity in the theta/gamma band was also shown to extend over similar areas of the probe, as was theta-gamma coupling during running movement (Sup. Fig. 3 and 4). Apart from arbitrary frequency bands, spectrogram analyses were performed using the Stockwell transform (Moukadem et al., 2015), which utilizes a sliding window to allow comparable resolution across frequencies (0-250 Hz in this case) for time-varying signals (Fig. 2C, 4C).

### 2.4 Ripple detection and propagation analysis

Before pursuing finer analysis on ripple propagation, it is important to note that NP 1.0 is equipped with 32 analog-digital converters (ADCs) multiplexing 12 channels each, which are sequentially sampled. This results in an inter-channel sampling offset of 33.33 μs which cumulates across channels within an ADC leading to an overall lag of 399.96 μs (∼ 0.4 ms) between first and the last channel of an ADC. It is important to correct these lags prior to ripple detection.

For ripple propagation detection, LFP was bandpass filtered in the range of 110-240 Hz using 4^th^ order Butterworth filter. The filtered trace was then Hilbert transformed to capture the ripple power which is then shown by its absolute value. Putative events were classified as ripples if the magnitude of the Hilbert transform exceeded a threshold of 5 × SD. Events shorter than 50 ms or longer than 200 ms were discarded. The ripple examples shown are from a dorsal tangential insertion (Fig. 2C-H). Ripple activity was prominent across more than a hundred channels showing the extended localization of ripple events. The given figure, however, shows every 4^th^ channel to avoid delays originating from the NP1.0 checkerboard pattern of the channels on the probe (Fig. 2E-H). For ripple propagation, quantification was done within each ripple event, using the start-time as the time at which the Hilbert envelope crossed 50% of its maximum value in each channel for that event (Fig. 2E, right panel); the time at which each channel reached maximum power, according to the Hilbert transform, was also included. To further extract the direction and speed of the propagation, a linear polynomial was fitted over the start times across all the subsequent channels. Note that the non-overlap between red line (fitted line over start time across channels) and dashed blue line (zero lag w.r.t the ripple onset at the initiation channel) shows that ripples propagate across channels.

### 2.5 Spike sorting, quality metrics

Spike sorting was done using Kilosort 2.5 (Fig. 4, (Pachitariu et al., 2016)). Standard cluster quality metrics were calculated to remove units of bad quality (Fig. 3E, Isolation distance > 10, refractory period violation (ISI) < 0.05%). All single-units were manually cured to search for over-clustered neuronal waveforms on Phy2 (Rossant et al., 2016). Once curing and quality metrics were applied, waveforms were locally recalculated by averaging the raw NP data at all spike times from this single-unit (up to 50 000 spikes, when possible, Fig. 5D). These recalculated waveforms should also include compensations for the slight offset correction of 2.78 μs between neighboring channels within the same analog-digital-converter, as previously done (Gehr et al., 2023; Sibille et al., 2022; 2024). This offset correction should be distinguished from the offset correction done in the LFP extraction for the ripple analysis (cf. above). In addition, to finalize the manual curing, a custom-written Graphical User Interface (GUI) was made to label the different single-units as good based on: 1 proper waveforms shape; 2 the absence of a prominent peak in the ±0.1 ms lag in its ACG; and 3 the absence of a peak in the ±0.3 ms lag in the CCG with all possible other combinations with other single-units.

Finally, an approximate distinction between putative excitatory versus putative inhibitory neurons was based on the shape of their waveforms at the channels of biggest amplitude (usually named best channel). To do so, we split waveforms at 0.42 ms of their peak-to-trough duration to distinguish between broad waveforms neurons (BW) and narrow waveforms neurons (NW), as a crude approximation of putative excitatory and inhibitory neurons respectively (cf. Sup. Fig. 5). According to a recent report using channel rhodopsin based optotagging (Valero et al., 2025), it is likely that our broad waveform neurons may also include Vip and Id2 inhibitory neurons which have larger waveforms.

### 2.6 Running and facial movements sampled at high frequency

Head-fixed mice were placed on a running band (Fig. 4A). For time alignment purposes, the different time triggered logic (TTL) pulses were all recorded on an extra NI-DAQ card (PXI-8381) connected to the NI-PXI recording system. These TTL included the 1 s long regular TTL pulse from the NP recording card, the camera frame triggers (at 150 to 200 Hz), and the analog signal from the running band (continuously recorded at 30 to 40 kHz). For all recordings, 1 camera (Basler acA1920-155c) was placed to monitor facial movements (Fig. 4B and Sup. Fig. 6, which can be improved to 3 to 5 cameras to monitor different sides of the face, and pupil of the animal). Illumination was done with IR lamps (ACL25416, Thorlab), equipped with a condenser Lens (Aspheric ACL25416U-B, Thorlab), positioned on the recorded side of the animal body, focused on the face. A separate computer was used to record the camera synchronously, which was externally triggered for each frame via a TTL pulse generated at 150-200 Hz by a dedicated locally made python script. Once recordings were finished, the movement data were extracted from the video using the singular value decomposition (SVD) from Facemap (Syeda et al., 2024) to plot vibrissae movement for each frame, allowing us to isolate moments of higher whisking activity versus inactive periods (among other facial movement, Sup. Fig. 6C). In addition, despite mice being head-fixed, regular run bouts could be extracted from the analogic run signal, which occurred less often (Fig. Sup. 6C: 2324 run events in 8 mice and 5817 whisking events in 14 mice), likely due to the lower sensitivity of band to the animal’s movements, which may reduce the detection efficiency of running events compared to un-constrained whisking events. Furthermore, whisking events were shorter in duration (Fig. Sup. 6C, averages FM = 300 ± 527 ms, n mice = 14, averages run = 967 ± 2067 ms, n mice = 8, p = 1.98*10^−159, Wilcoxon signed rank test). Putative run/whisker bouts were extracted based on a threshold of 3 X SD of the movement trace. To account for artefact-related detections, bouts shorter than 10 ms were removed for further analysis. There was an expected increase in theta power during the movement bouts (Fig. 4C). Next, we analyzed the hippocampal neuronal correlates of whisking (Fig. 4E) and running behavior (Fig. 4F). Spiking activity at the level of single-units was assessed 500 ms before and after the onset and offset of movement trials. We found an overall increase and decrease in the spiking activity after the movement onset and offset respectively for both whisking and running. To quantify the proportions of neurons contributing to the change in the firing, a threshold of ± 3 X SD of spiking activity after the movement onset/offset was used.

### 2.7 Functional synaptic connectivity detection and quantification

Connectivity in the hippocampus between different neurons was addressed using cross-correlation analysis (CCG), using pycorrelat (Laurence et al. Optics Letters(2006)) similarly to what was done in the visual system (Gehr et al., 2023; Sibille, Gehr, Teh, et al., 2022) and in the hippocampus (English et al., 2017; Mizuseki & Buzsáki, 2013).

Neurons having less than 2000 spikes were discarded from CCG analysis. CCG was calculated with a bin size of 0.1 ms. A copy of the spike trains, referred to as spike jitters was estimated by randomizing every spike within a 10 ms window, which was then subtracted from the raw CCG. This step, referred to as CCG-jitter correction, removes the putative common input biases which may not be related to mono-synaptic delays. Excitatory contacts were identified by a prominent peak in monosynaptic delays (0.5 – 3 ms), as done previously (Liew et al., 2021a). Putative inhibitory connections were not considered due to the difficulty of detection in the corresponding CCGs. For every pair, different properties were extracted such as the relative distance between the pre-synaptic and post-synaptic neurons, computed by Euclidean distance between their respective best channels (Fig. 5F). Because we observed cases of multiple contacts emerging from a single neuron toward several other neurons, or inversely multiple neurons converging toward a neuron, we calculated divergence and convergence of synaptic contacts (Fig. 5F). To estimate the synaptic efficacy of each pair (Fig. 5F), the baseline of the CCG (−5 ms to 0 ms) was subtracted from the area of their CCG peak (0.5 ms to 3 ms). To estimate synaptic efficacy and contribution, it is important to avoid using jitter-corrected CCGs, because they can contain negative values which will lead to an underestimation of efficacy values. Synaptic efficacy was defined as the ratio of the peak area (baseline corrected) and the pre-synaptic spike number, while contribution was defined as the ratio of the peak areas (baseline corrected) to the post-synaptic number for the post-to-pre recalculated CCG.

## Data and code availability

Data and code are available on request.

## 3. Results

### Tangential insertions maximize spatial sampling yields in the hippocampus

First, to illustrate the advantages of tangential hippocampal insertions, we compared the localization of channels within the hippocampal formation across different insertion types (Fig. 1B). Although qualitatively we observed more channels in the hippocampus as a whole, in CA1, and in CA1-3 (which we denote as CAX) in the tangential insertions than in the more orthogonal ones, the difference was only statistically significant for the number of channels across the whole hippocampus in tangential ventral (371 ± 23, n = 3) versus orthogonal insertions (229 ± 29, n = 3) (p = 0.049, Wilcoxon rank-sum), and for the number of CA1 channels in tangential dorsal (200 ± 19, n = 8) versus orthogonal insertions (112 ± 39, n = 3) (p = 0.014, Wilcoxon rank-sum). Given the very small number of orthogonal insertions (n = 3) in our dataset, such comparisons are statistically under-powered, but the trend for higher number of hippocampal channels in the tangential insertions is clear (also in comparison to other datasets from more purely orthogonal recordings, see discussion), except for DG.

Second, we characterized the LFP profiles across all the channels, comparing the number of channels with supra-threshold ripple, gamma, and theta band activity across each insertion type (Fig. 2B). Despite the low statistical power mentioned above, the number of channels across which we could detect ripples (Fig. 2B, top) was significantly larger during tangential compared to orthogonal insertions (dorsal (120 ± 39, n = 7) vs orthogonal (24 ± 1, n = 3): p = 0.017; ventral (73 ± 45, n = 3) vs orthogonal: p = 0.049, Wilcoxon rank-sum; note one dorsal recording had to be excluded due to lack of ripples). Because of our much higher spatial sampling within the hippocampus, we are able to track individual ripples across large swaths of the hippocampus using this method (Fig. 2). We also found a broader coverage of oscillatory gamma and theta activity in tangential insertions compared to orthogonal insertions. The number of channels encompassing higher theta power was significantly enhanced in both kinds of tangential insertions (dorsal (177 ± 42, n = 8) vs orthogonal (95 ± 29, n = 3): p = 0.014; ventral (176 ± 24, n = 3) vs orthogonal: p = 0.049, Wilcoxon rank-sum), while the number of channels encompassing higher gamma power was enhanced for dorsal tangential insertions (dorsal (220 ± 48, n = 8) vs orthogonal (65 ± 14, n = 3): p = 0.014) and decreased for ventral tangential insertions (ventral (37 ± 5, n = 3) vs orthogonal: p = 0.049, Wilcoxon rank-sum).

Third, we investigated hippocampal single-unit yield across all insertion types. Maximal cell yield achieved in a single recording in dorsal tangential insertion reached about 710 units compared to 384 and 168 units (Supplementary Tab 1) in ventral tangential and orthogonal insertions respectively. Qualitatively, it seems that the yields are highest in the dorsal tangential insertions, but again the low statistical power limits the certainty of such comparisons: we did not observe any group level significance (Fig. 3D) in the cell reach across different insertion types (dorsal (313 ± 218, n=8)/ventral (174 ± 182, n=3): p = 0.497; dorsal/orthogonal (125 ± 57, n = 3): p = 0.376; ventral/orthogonal: p = 1.000, Wilcoxon rank-sum).

Fourth, we compared several measures of recording quality between different insertion types. Similar to what has been described for tangential insertions in the superior colliculus (Sibille, Gehr, Benichov, et al., 2022), we report a significantly lower drift (Fig. 3F left panel) in dorsal hippocampal tangential insertions (12.01 ± 7 µm, n = 8) compared to orthogonal (15.75 ± 10.8 µm, n = 3; p = 3.49*10^−16; Wilcoxon rank-sum test) or ventral tangential insertions (15.35 ± 11 µm, n = 3; p = 1.08 *10^−14; Wilcoxon rank-sum test). ISI and Isolation distance were also significantly improved in tangential insertions compared to orthogonal (Fig. 3F middle and right panels; comparisons for dorsal vs ventral, dorsal vs orthogonal, and ventral vs orthogonal: ISI, p = 0.28/0.006/0.03, Isolation distance, p = 4.07*10^−09/0.001/4.36*10^−06, ; Wilcoxon rank sum tests, n = 2505/523/375 neurons from n mice = 8/3/3 for dorsal, ventral, and orthogonal, respectively).

We next investigated the possibility to combine our insertions with behavioral tracking as classically done with head-fixed awake experimentation. We found a profound temporal relationship between neural activity and whisking compared to running bouts for both onset and offset of movements (Fig. 4G). We note that the larger delays in onset and offset responses to running bouts versus whisking may just reflect the delays that exist between the bodily movement in the mouse and the actual band movement during run. This delay may also partially underlie the earlier emergence of theta oscillations in the hippocampal LFP compared to the running band signal (Fig. 4C). Taking the paw/breast motions into account (which can be tracked from the facial video, Fig. 4B) could potentially resolve this issue but is unfortunately confounded by other movements such as respiration (cf. Sup. Fig. 6).

The kinetics of recruitment in the beginning and end of whisking bouts (Onset & Offset, for Facemap and run) showed very small differences in BW versus NW neurons for vibrissae movements (av. onset = 22.9 ± 79/15.1 ± 63 ms for BW and NW respectively; p = 0.5,n = 1649/368 units, n = 14 mice; av. offset = -16 ± 93/ 16 ± 59 ms for BW and NW respectively; p = 2.51 * 10^−8, n = 1465/302 units, n = 14 mice, Wilcoxon rank sum). During a run bout, in agreement with the delay observed on theta oscillations, we report similar delay at the level of single-units (av. onset = 76 ± 98/79 ± 92 ms for BW and NW respectively.; p = 0.63, av. offset = -40 ± 94 / -39 ± 92 ms for BW and NW respectively., p = 0.88, n = 106/382 units, n mice = 8, Wilcoxon rank sum).

Fifth, we compared the functional connectivity by quantifying the excitatory synaptic contacts across all insertion types, and found no statistical significance (Fig. 5F, average pairs detected = 88 ± 97/33 ± 29/56 ± 44, dorsal/ ventral/orthogonal respectively, p = 0.54/0.83/0.27, n mice = 8/3/3, Wilcoxon rank sum). Despite not reaching group level significance, maximal number of synaptic connections detected in a single recording in the dorsal tangential insertion reached up to 243 as opposed to 65 and 98 in ventral tangential and orthogonal insertion types respectively. Overall, as in sensory (English et al., 2017; Valero et al., 2025). In addition, unlike previous reports showing mostly excitatory neurons to inhibitory neurons (English et al., 2017; Mizuseki & Buzsáki, 2013), we observed regularly excitatory neurons to excitatory neurons connections. These two types of excitatory connections exhibit different features depending on whether their post-synaptic target is BW or NW neurons. The average peak time for BW targeted connections was significantly higher (2.3 ± 1.19 ms, n = 248) compared to the NW (1.63 ± 0.66 ms, n = 730) connections (p = 8.16*10^−18, Wilcoxon rank-sum). Similarly, average efficacy (eq. to the probability of spike transmission) was higher for NW targeted connections (5.72 ± 5.6%) compared to BW (3.4 ± 3.39%) connections (p = 4.21*10^−15, Wilcoxon rank sum) while the contribution (eq. to the proportion of post-synaptic spikes locked to pre-synaptic inputs) was modestly lower for NW connections (2.06 ± 2.4%) compared to BW (2.80 ± 3.8%) connections (p = 0.0041, Wilcoxon rank sum). Furthermore, we did not observe an overall difference in the distance of the excitatory connections onto NW or BW neurons. Interestingly, we did observe unusually long connection distances beyond 400 µm for both connection types corresponding to inter-region connections (Fig. 5D & F, p = 0.96, avg. BW/NW = 139 ± 182/129 ± 148 µm). We found these long-range connections specifically in the recordings encompassing many channels within the pyramidal layer (3 out of 1with multiple hippocampal regions showed prominence of, as previously observed (English et al., 2017).

## 4 Discussion

In this paper, we illustrate a novel technique of inserting a single NP tangentially along the hippocampal antero-posterior axis to maximize the spatial coverage, in head-fixed mice (Fig. 1). Tangential insertions have been done previously in the retinotectal (Sibille, Gehr, Benichov, et al., 2022) and thalamo-cortical (Sibille et al., 2024) visual systems in mice, but the approach has not been tested in the hippocampus.

We hereby highlight the key advantages of this technique including: increased channel yield from hippocampus, both from dorsal (whole hippocampus) and ventral (CA1) tangential insertions (Fig. 1), allowing for the tracking of LFP patterns with high spatiotemporal precision, as illustrated by ripples (Fig. 2); the ability to record a large number of neurons from stratum pyramidal over an extended anatomical range (Fig. 3); together with more classical behavioral tracking (Fig. 4); the potential of capturing synaptic connections across the extended hippocampal spatial domain (Fig. 5). Consequently, this approach opens new avenues in analyzing possible relationships between anatomical patterns, connectivity, LFP, and behavioral variables across the longitudinal axis of hippocampus.

We show via extensive camera tracking correlations between hippocampal neuronal activity and behaviors such as whisking or running (Dudok et al., 2021; Rios et al., 2025). It is important to highlight that vibrissae movements are rapid, occurring sometimes within 100 ms, hence extracting facial movements below 20 Hz sampling rate would result in an overestimation of resting state (Sup. Fig. 6). The animals in the current set-up were well habituated, therefore alternating between stable silent and mobile run periods. Running periods, however, were classified based on the output from the treadmill, which was preceded by actual bodily movements in mice, thereby slightly underestimating the actual run bout durations. This is exemplified by earlier onset of theta oscillations compared to the initiation of run bout (Fig. 4).

While imaging techniques have also been developed in parallel to record the activity of large numbers of neurons from cortical (Abdelfattah et al., 2022; Kim et al., 2016; Stringer et al., 2019), and subcortical regions (Sun et al., 2025), they still lack temporal resolution due to their reliance on voltage- or calcium-sensors and/or the tradeoff between scanning speed and image volume. Furthermore, such studies entail elaborate and more expensive setups, which may not be necessary for most experiments requiring simpler readout. Because NP recordings can also be applied to humans (Chung et al., 2022, 2025; Coughlin et al., 2023; Paulk et al., 2022), knowledge gained from recordings in rodents (as in our case) and other species (e.g. (Town et al., 2023; Trautmann et al., 2025)) can in principle be directly compared to human data, if anatomical differences are considered.

The hippocampus exhibits anatomical and functional gradients across its longitudinal axis and tangential insertion of high-density NP across this axis provides a unique opportunity to capture the neural activity with higher spatial resolution. This allows the study of network dynamics including sharp wave ripples which may vary systematically across the axis. Propagation of sharp wave ripples across the longitudinal axis has previously been studied, the sampling however was performed at discrete spatial points across the axis limiting the spatial resolution (De Filippo & Schmitz, 2023; Patel et al., 2013).

In principle, our approach could be extended to other probe types and recording conditions. Several approaches have been adapted to maximize the neuronal yield for e.g. insertion of multiple probes in parallel (Durand et al., 2023; Luo et al., 2020), multi-shank (Steinmetz et al., 2021), or of high-density probes (Angotzi et al., 2025; Paleologos et al., 2025; Ye et al., 2025). These approaches have been primarily designed to maximize the neural outputs from various brain regions and are not necessarily hippocampal-specific. Usage of multiple probes, however, can be technically challenging due to spatial constraints making physical implementation difficult. Multi-shank probes offer an advantage over single shank probes, but they still lack the high spatial sampling our approach offers across the longitudinal axis of the hippocampus.

Previously, Swanson et al., 2025 used an angular approach with silicon probes. They inserted silicon probes at an angle of 60° within the coronal plane since their goal was to simultaneously target retrosplenial cortex and dorsal hippocampus. Consequently, their coverage within the pyramidal layer was relatively limited, with just 8-10 channels (Swanson et al., 2025) compared to our dorsal tangential insertions encompassing more than 100 channels within the pyramidal layer (Fig. 2D-E-F). Application of orthogonal and ventral tangential insertions seem feasible in freely moving mice, as recently published using a 60° angle from the posterior midline (Swanson et al., 2025). This may be challenging for dorsal tangential insertions in mice as the probe is inserted from the anterior midline at an angle of approximately 30° from the midline plane rendering the implant accessible for the animal, potentially causing mechanical disruption. In addition, this positioning of the NP at a flat angle will make the implant very large and extended toward the frontal part of the head, which may also hinder the head movement in the animals. Previously, feasibility of tangential insertions in the cortex of rats in chronic settings was shown to be possible at an angle of 65° (Mimica et al., 2023) suggesting that all tangential hippocampal insertions could be done in rats. Tangential insertions can also be combined with a head-mounted multi-camera system for behavioral tracking in freely moving mice (González-Rueda et al., 2024; Meyer et al., 2018).

In the current study, we have explored only the acute tangential insertions in head-fixed conditions and have shown that they can substantially increase the sampling within the hippocampus compared to more orthogonal insertions, enabling us to capture local field potential dynamics across a larger extent of the longitudinal axis. This facilitated the detection of ripples in up to 130 channels corresponding to about 1.3 mm of tissue, allowing the investigation of ripple dynamics including propagation and coordination. In terms of cell and synaptic yield, we did not observe a statistical significance between tangential and orthogonal insertion types. However, maximal cells and synaptic yield were higher for dorsal tangential insertion compared to ventral tangential or orthogonal insertions. It is to be noted that orthogonal insertions were performed at an angle of 45° from both the sagittal and the coronal plane. Although we did not observe any statistical difference in the cell yield between orthogonal and tangential insertions, our angled approach collectively across all insertion types markedly increased cell yield from hippocampus compared to purely vertical approached done previously (Jun et al., 2017).

Additionally, we observed diverse connectivity profiles including locally connected circuits and more broadly connected networks (Fig. 5D-E). The lack of statistical significance could be attributed to the limited sampling size and higher variability, possibly stemming from the sensitivity of the placement of the probe right next to the pyramidal layer(s) of the hippocampus. Furthermore, our findings are in line with previous studies regarding the prominence of excitatory mono-synaptic connections toward putative inhibitory neurons (Fig. 5, English et al., 2017; Mizuseki & Buzsáki, 2013). Given the methodological limitations of detecting connectivity and its inconsistency with patch-clamp based estimates (Bertagna et al., 2024; Ghanbari et al., 2020; Liew et al., 2021b; Sammons et al., 2025), it remains to be understood whether these outcomes truly reflect the physiological connectivity.

## Acknowledgments

We would like to acknowledge the entire Schmitz lab for providing support: Roberto de Filippo who wrote the license, Antje Fortströer, Anke Schönherr, and Friedrich Johenning who supported the underlying administrative work.

## Author Contributions

Conception and design: J. S.

Acquisition of data: A. K., J. S.

Analysis and interpretation of data: A. K., V. E., J. T., J. S.

Drafting or revising the article: A. K., J. K., V. E., J. T., D. S., J. S.

## Supplementary figures

**Supplementary Figure 1:**
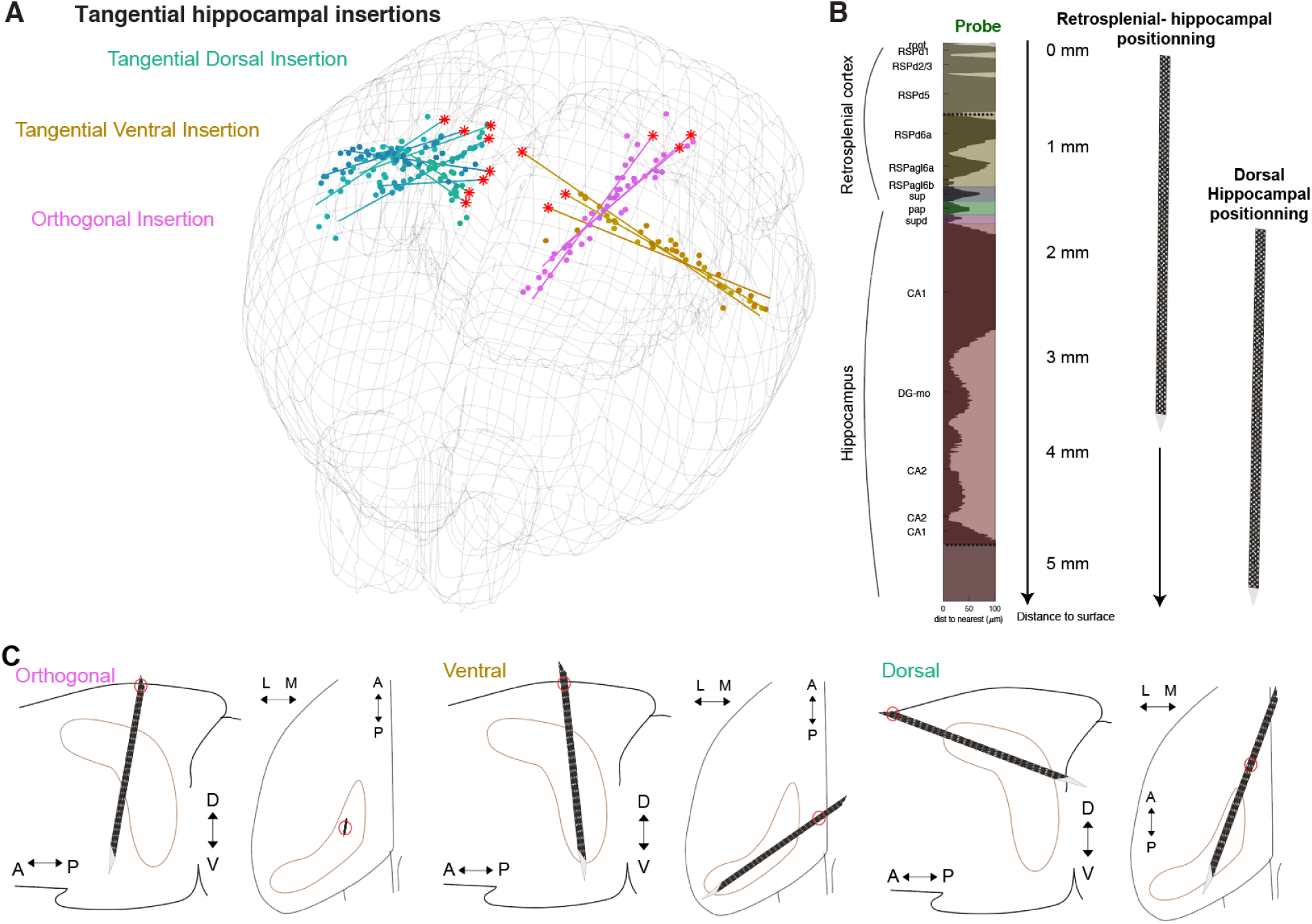
Dorsal and ventral tangential insertions require deep insertions to reach the hippocampus: **A.** Schematic of tangential (dorsal and ventral) and orthogonal insertions (n = 14 mice). **B.** The 5 mm track of the dorsal tangential insertion spans several bran regions (left), where two different positions can be achieved: a Retrosplenial-hippocampal positioning (middle), or a full hippocampal positioning (right). **C.** Sagittal and interaural planes represent the position of the insertion for orthogonal (left), ventral (middle), and dorsal (right) insertions. Note that comparatively the point of entering the tissue (red circle) is quite variable from one insertion to the other (interaural view).

**Supplementary Figure 2:**
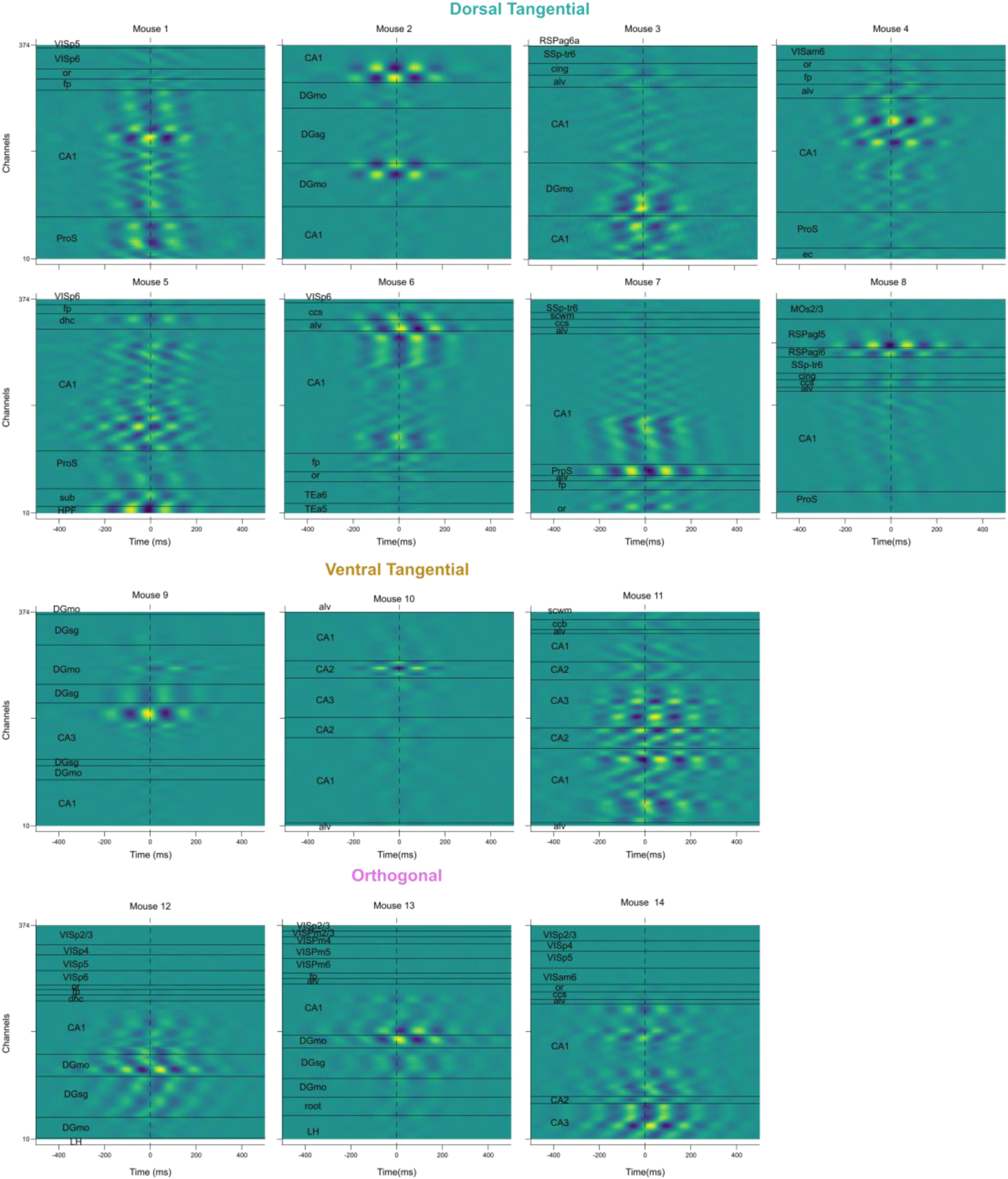
Single recording CSD aligned to theta trough. CSD calculated along each probe aligned to theta trough. Timestamps for theta trough were taken from the channel with the highest theta power for each recording. List of anatomical abbreviations can be found in supplementary table 2.

**Supplementary Figure 3:**
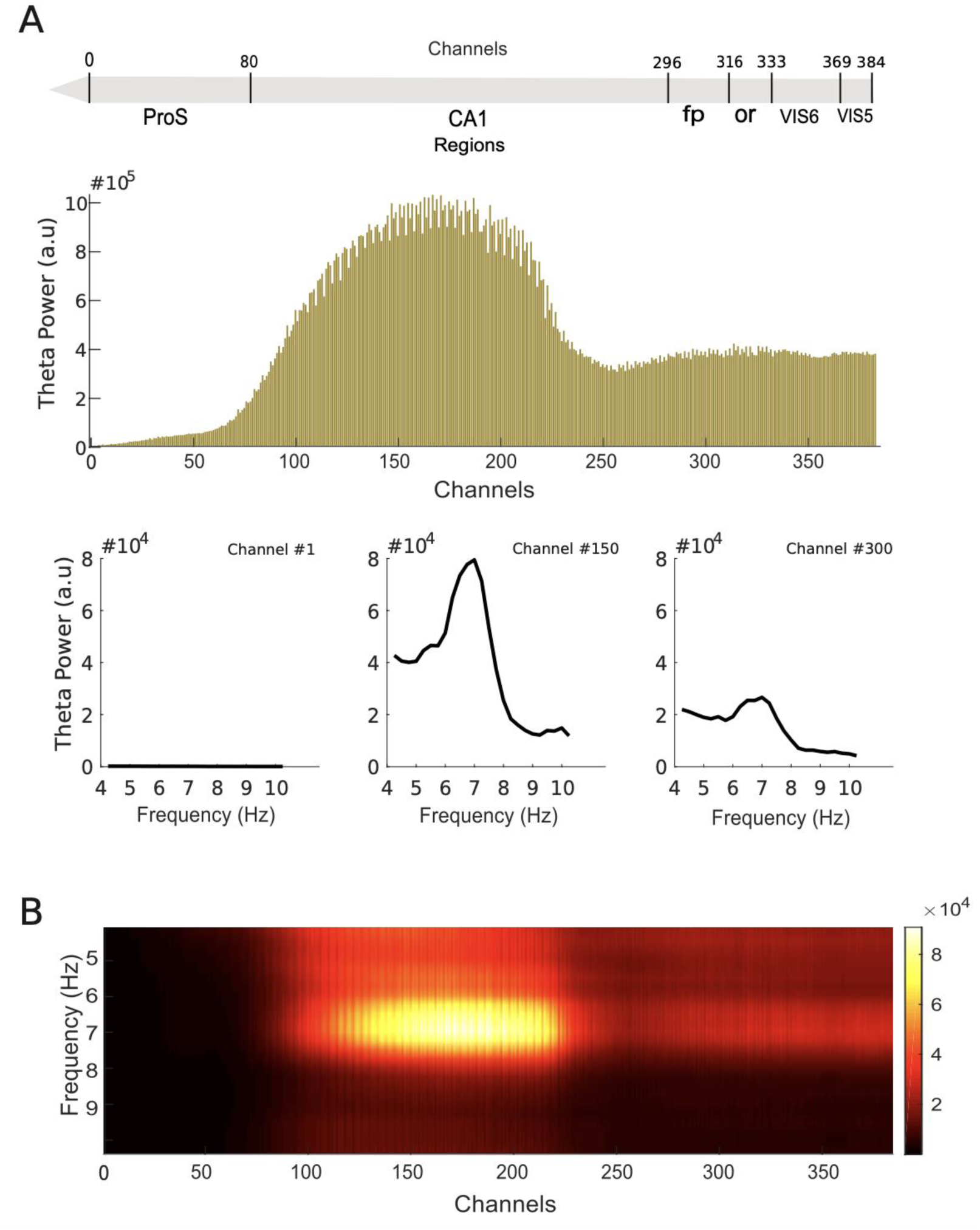
Differential theta power distribution across NP during a 20 s long running movement epoch. **A.** Anatomical coverage across channels in a representative recording (Top). ProS - Presubiculum, CA1 - Cornu Ammonis 1 (hippocampal CA1 pyramidal layer), fp – corpus callosum posterior forceps, or - Stratum oriens, VIS6 - Primary visual cortex, layer 6, VIS5 - Primary visual cortex, layer 5. Histogram of summed theta-band power (4–10 Hz) computed using FFT across all recording channels during the 20 s running epoch (Middle). PSD traces from three representative channels in deep (channel 1), middle (channel 150) and superficial (channel 300) locations of the probe (Bottom). **B.** Power frequency representation across the entire probe. The heatmap shows power as a function of frequency (y-axis) and channel depth (x-axis), demonstrating spatially localized theta-band activity.

**Supplementary Figure 4:**
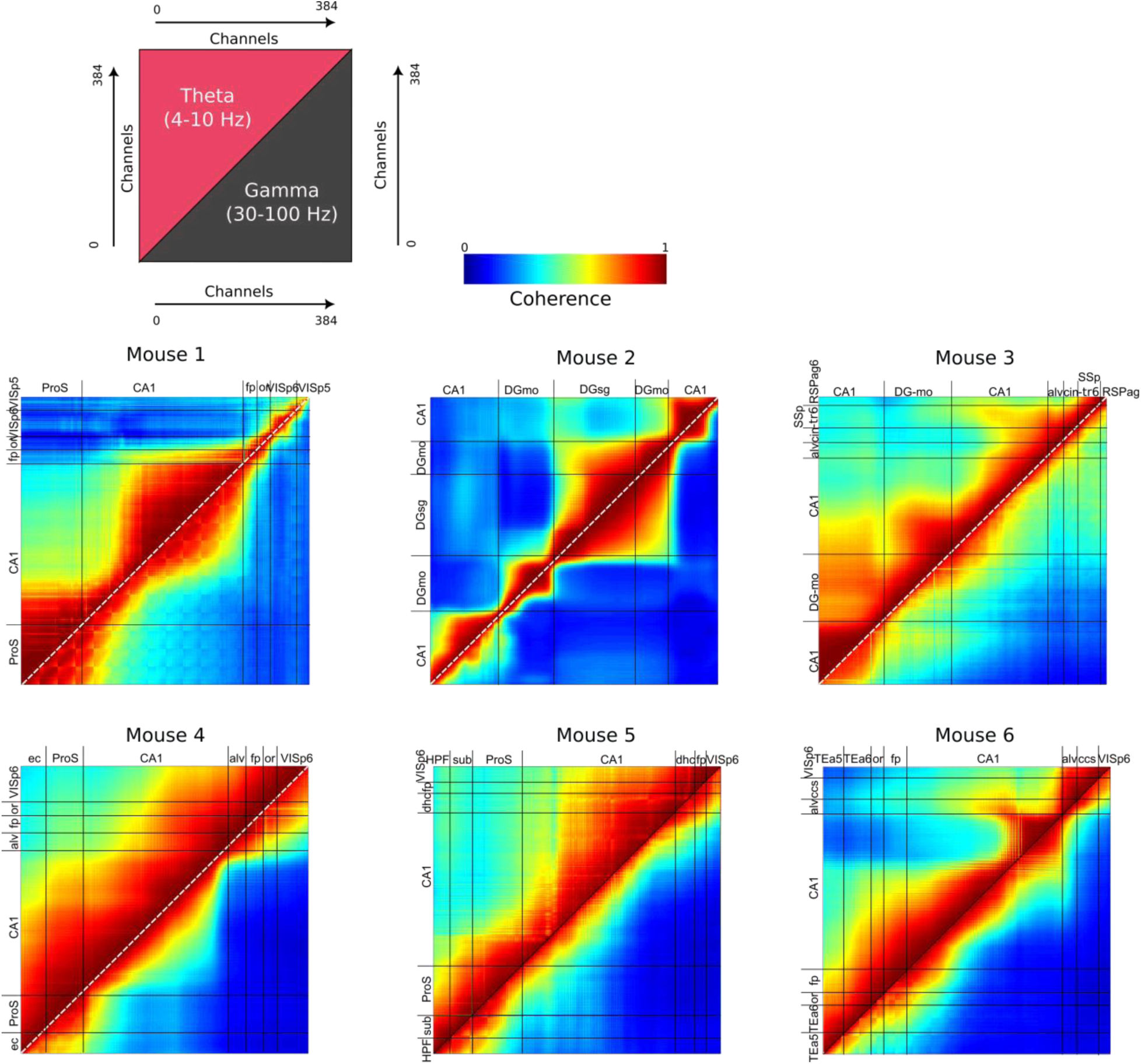
Movement-induced theta and gamma power increase. Scheme of the coherence quantification arranged along the probe channels (Top). All heatmaps show coherence along the 384 recording channels, from six different dorsal tangential recordings. Colors indicate the magnitude of squared coherence between two channels during movement, splitting theta (top-triangle) from gamma (bottom triangle) range. Brain regions are indicated in the top and left sides for each recording.

**Supplementary Figure 5:**
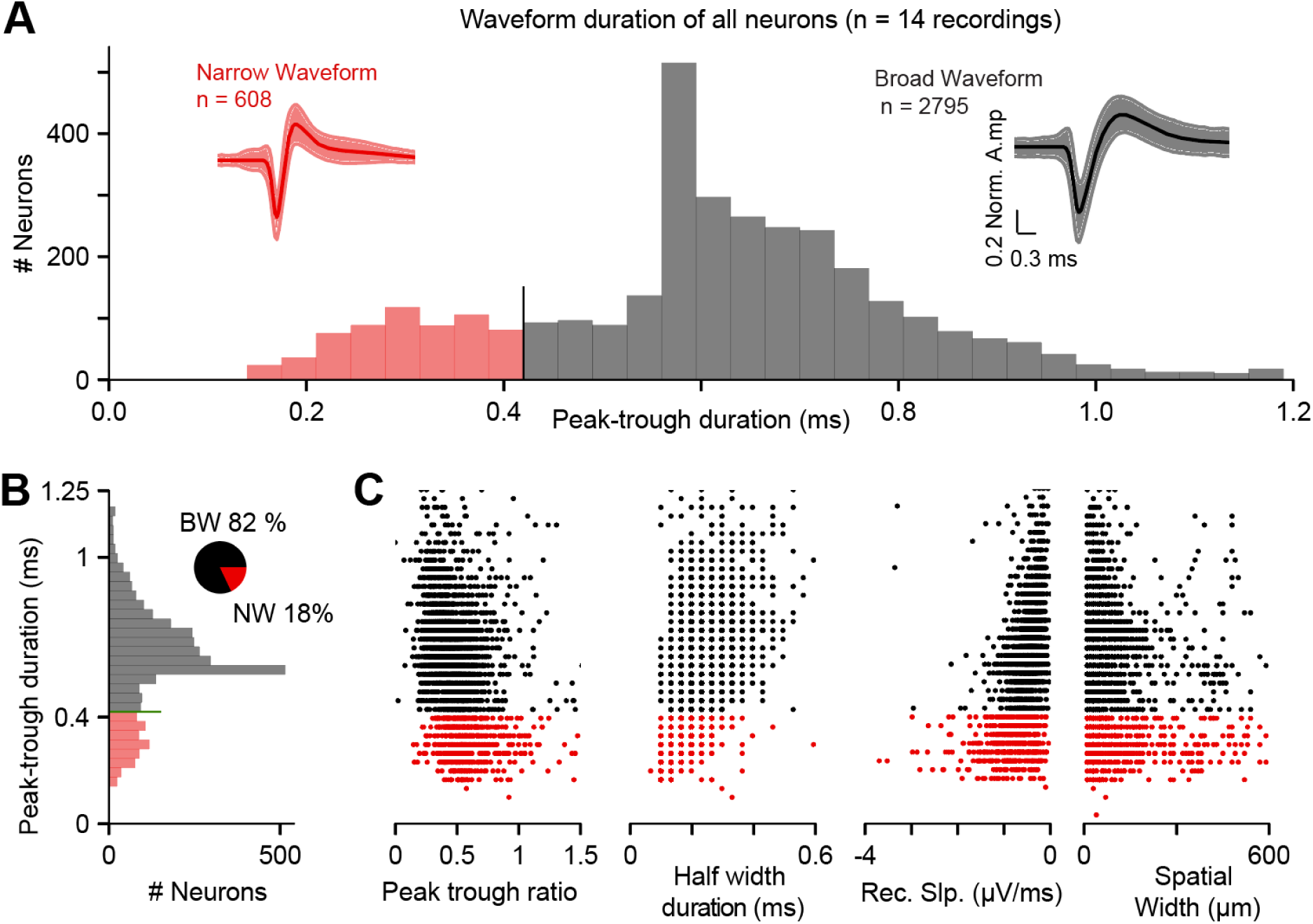
Narrow versus broad waveforms neurons in the hippocampus: an approximation to extract excitatory from inhibitory neurons. **A.** Typical peak-to-trough duration used to separate narrow waveforms (NW, red) from broad waveform (BW, black) neurons. Insets show the averages and std of the normalized waveforms from all neurons; scale bar applies to both inset identically. **B.** Peak-to-trough duration plotted vertically, aligned to other waveforms properties. **C.** Quantifications in NW versus BW neurons in the entire dataset; each dot represents either a BW (black) or NW (red) neuron. From left to right: peak-trough ratio, half width duration, recovery slope, and waveforms’ spatial width; all sharing the same y-axis as in B. Note that the 30 kHz sampling rate of NP give these incontiguous lines rendering in the different panels.

**Supplementary Figure 6:**
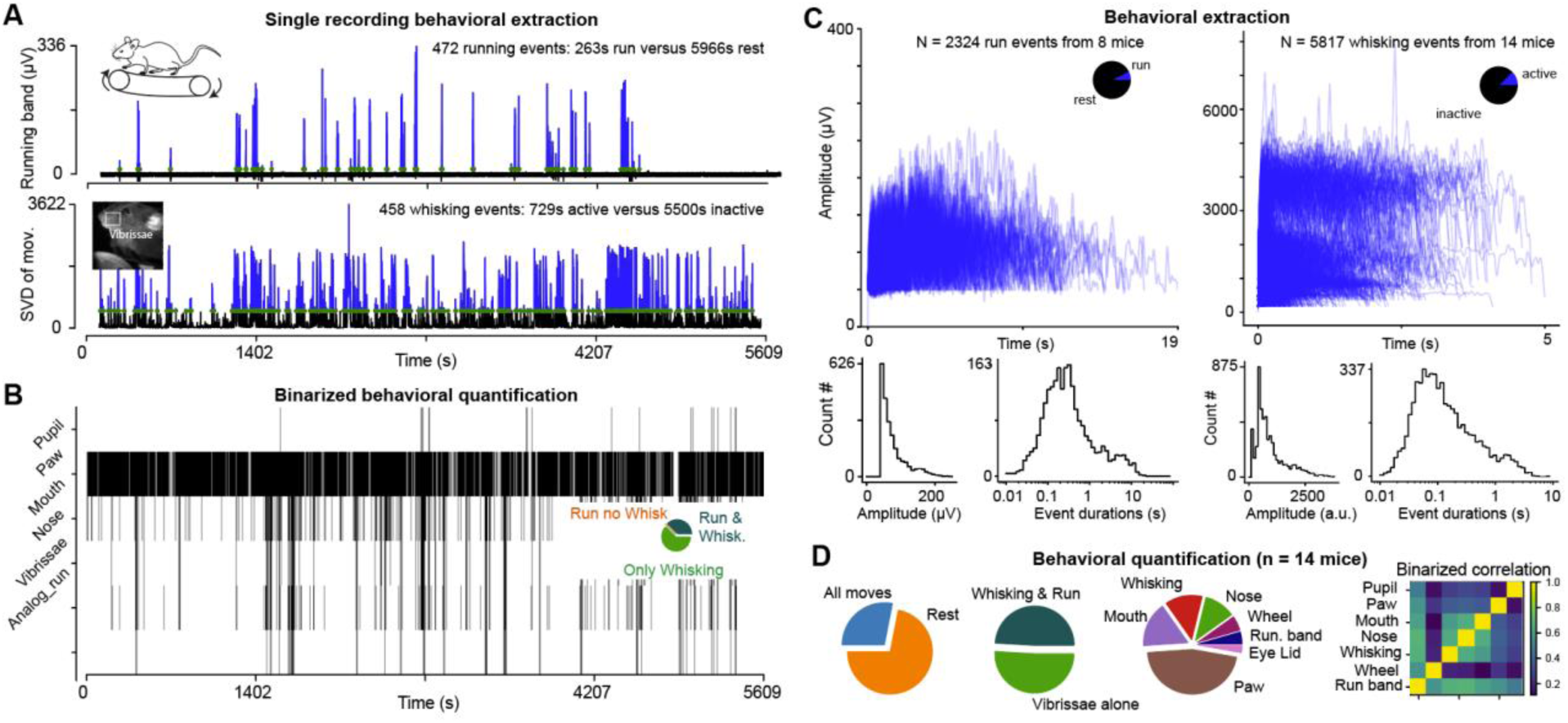
Spontaneous behavior (run and whisking) monitored at high frequency. **A.** Representative recording showing outputs from the running band (top) and singular value decomposition (SVD) signal from the facial movements (bottom, extracted with Facemap). Blue lines indicate the signal above the threshold selected for further analysis; black lines are sub-threshold; green dots indicate the point of threshold crossings. Both have a schematic inset of recording methods (left). **B.** Binarized overlayed behavior over time from the different parts of the video (top) and from the running band (bottom). Note the quantification in this animal of the proportion of synchronic run and whisking periods versus periods of whisking alone. **C.** Overlay of all extracted run bouts in all recordings (top); the pie plot indicates the proportion of active (“run”) and passive (“rest”) periods in the entire recording duration. Quantifications of its amplitude (bottom left) and duration properties on a logscale axis (bottom right). **D.** Dataset quantifications of time spent active (“All moves”) versus rest (left); of the period with whisking and run versus whisking alone (middle left), of the periods related to different facial movements (middle right) and their respective correlations (right). Note that as expected, mouth, nose and whisking periods are strongly correlated.

**Supplementary Tab 1:** Table summarizing recording wise channels in the hippocampus, CA1, cell yield, synaptic yield, channels with enhanced power in theta and gamma range, and number of channels with ripple activity.

| Mouse | Insertion | Ch. In Hippo | Ch. In CA1 | Cell yield (n) | Synaptic yield (n) | Theta (ch) | Gamma (ch) | Ripple (ch) |
| --- | --- | --- | --- | --- | --- | --- | --- | --- |
| 12 | Orthogonal | 243 | 91 | 61 | 10 | 67 | 78 | 25 |
| 13 | Orthogonal | 196 | 87 | 168 | 98 | 125 | 66 | 23 |
| 14 | Orthogonal | 248 | 157 | 146 | 61 | 93 | 50 | 25 |
| 9 | Ventral | 384 | 88 | 384 | 65 | 148 | 43 | 45 |
| 10 | Ventral | 384 | 227 | 71 | 5 | 99 | 73 | 125 |
| 11 | Ventral | 344 | 174 | 68 | 29 | 191 | 35 | 50 |
| 7 | Dorsal | 261 | 222 | 59 | 0 | 156 | 201 | 100 |
| 8 | Dorsal | 224 | 171 | 135 | 4 | 152 | 237 | 120 |
| 1 | Dorsal | 296 | 216 | 277 | 79 | 132 | 319 | 49 |
| 2 | Dorsal | 384 | 172 | 294 | 116 | 232 | 153 | 171 |
| 3 | Dorsal | 323 | 212 | 525 | 222 | 234 | 228 | no ripple |
| 4 | Dorsal | 268 | 194 | 364 | 243 | 212 | 194 | 129 |
| 5 | Dorsal | 349 | 207 | 710 | 30 | 147 | 199 | 151 |
| 6 | Dorsal | 227 | 208 | 141 | 16 | 152 | 226 | 121 |

**Supplementary Tab 2:** List of anatomical abbreviations.

|  |  |  |  |
| --- | --- | --- | --- |
| alv | alveus of the hippocampus | LH | Lateral habenula |
| CA1/2/3 | cornu ammonis 1/2/3 | MOs2/3 | Secondary motor area, layer 2/3 |
| ccb | corpus callosum, body | or | optic radiation |
| ccs | corpus callosum, splenium | ProS | Pro-subiculum |
| cing | cingulum bundle | root | root |
| DG-mo | Dentate gyrus, molecular layer | RSPagl5,6 | Retrosplenial area, lateral agranular part, layer 5,6 |
| DG-po | Dentate gyrus, polymorph layer | scwm | supra-callosal cerebral white matter |
| DG-sg | Dentate gyrus, granule cell layer | SSp-tr6 | Primary somatosensory area, trunk, layer 6 |
| dhc | dorsal hippocampal commissure | Sub | Subiculum |
| ec | external capsule | TEa5,6 | Temporal association areas, layer 5,6 |
| Ent-6a | entorhinal area, lateral part, layer 6a | VISp2/3,5,6 | primary visual area, layer 2/3 |
| fp | corpus callosum posterior forceps | VISam6 | anteromedialvisual cortex, layer, 6 |
| HPF | hippocampal formation | VISpm2/3,4,5,6 | posteromedialvisual cortex, layer, 6,5,4,2/3 |

